# Network and behavioral signatures of postpartum behavioral deficits improve with a neurosteroid-based treatment

**DOI:** 10.64898/2026.09.25.754460

**Authors:** Lifen Htet, Grant Weiss, Kaitlyn E. Dorst, Sarah Gray, Mike Lewis, Jamie Maguire

## Abstract

Postpartum depression (PPD) impacts nearly 20% of mothers and a much larger percentage (up to 75%) suffer from postpartum blues. Despite this high incidence, we still have an incomplete understanding of the underlying neurobiology of PPD. Deficits in neurosteroid signaling has been implicated in the vulnerability to postpartum behavioral deficits and mice with reduced expression of δ subunit-containing GABA_A_ receptors (GABA_A_Rs) (*Gabrd* ^-/-^ or *Gabrd* ^+/-^), the predominant site of action for 5α-reduced neurosteroids, exhibit behavioral deficits restricted to the postpartum period. Allopregnanolone and allopregnanolone analogs (SGE-516) can improve postpartum behavioral outcomes and their ability to modulate oscillations in the basolateral amygdala (BLA) involved in emotional processing has been proposed to contribute to their antidepressant effects. Here we demonstrate that the behavioral differences in postpartum wild type, *Gabrd*^*+/-*^, and SGE-516-treated *Gabrd*^*+/-*^ dams is largely driven by theta oscillations in the BLA. Previous studies have linked theta oscillations in the BLA to altered behavioral responding in response to stress or threat. Interestingly, here we demonstrate that *Gabrd*^*+/-*^ dams exhibit a failure to properly respond to their own pup calls, with reduced theta oscillations that correlates with deficits in maternal approach behavior both of which are restored with SGE-516 treatment. Further, unbiased MoSeq analysis identified altered behavioral syllable expression in *Gabrd*^*+/-*^dams in response to their pup calls compared to wild type dams that are restored with SGE-516 treatment. These findings demonstrate network and behavioral signatures of postpartum behavioral deficits and characterize the effects associated with neurosteroid-based treatment response.

## Introduction

Postpartum Depression (PPD) is a serious mental illness that impacts nearly 20% of mothers shortly after giving birth, with substantial higher rates in developing countries and among minoritized communities [1]. Based on the timing of symptom presentation, fluctuations in ovarian hormones have been implicated in the pathophysiology of PPD. Lower levels of allopregnanolone, a metabolite of progesterone, have also been implicated in PPD [2] and shown to predict postpartum depression [3] (for review see [4]).

Allopregnanolone is a potent positive allosteric modulator (PAM) of GABA_A_ receptors (GABA_A_Rs), particularly extrasynaptic, δ subunit-containing GABA_A_Rs [5, 6]. Our previous research found that mice that exhibit reduced expression of the GABA_A_R δ subunit, *Gabrd* ^+/-^and *Gabrd* ^-/-^ mice, exhibit abnormal postpartum behaviors restricted to the postpartum period, including increased stress-induced helplessness and deficits in maternal care [7]. Further, we demonstrated that treatment with a synthetic allopregnanolone analog, SGE-516, improves abnormal postpartum behaviors in two independent preclinical models which exhibit postpartum behavioral deficits (Melon and Maguire, 2018). Similar synthetic allopregnanolone analogs received FDA approval for the treatment of postpartum depression, brexanolone [8] and zuranolone [9].

The mechanisms mediating the therapeutic effects of allopregnanolone analogs are still not completely understood, but are thought to involve their ability to act as PAMs at GABA_A_Rs and modulate network states involved in emotional processing, including the basolateral amygdala (BLA) [10]. Amygdalar dysfunction has been implicated across psychiatric illnesses [11, 12], including postpartum depression [13]. Previous research suggests that the rhythmic synchronization within and between the BLA and PFC circuits is required for behavioral responses related to stress and threat [10, 14]. Specifically, distinct oscillatory states such as low 2-5 Hz and 6-12 Hz high theta bands are associated with fear and safety behavioral states, respectively, with a significant difference in the ratio of these frequencies [14, 15]. Our laboratory previously demonstrated the ability of allopregnanolone modulate BLA oscillations, facilitating pro-safety oscillations (6-12 Hz) through actions on δ-containing GABA_A_ receptors, the effects of which were similar in both rodents and humans [10].

Here we investigate whether the ability of allopregnanolone analogs to improve postpartum behavioral outcomes involves the ability of these compounds to modulate oscillations in the BLA related to emotional processing. We utilized a preclinical model with reduced expression of the GABA_A_R δ subunit, *Gabrd*^*+/-*^ dams, to interrogate the network and behavioral states contributing to the postpartum behavioral deficits and disrupted maternal care. We demonstrate that *Gabrd*^*+/-*^ dams exhibit a failure to properly respond to their own pup calls both at the level of behavioral and network states. We demonstrate that altered BLA network states are associated with deficits in maternal care, which is improved with SGE-516 treatment. Further, *Gabrd*^*+/-*^ dams exhibit altered syllable expression measured by MoSeq compared to wild type dams, many of which are restored with SGE-516 treatment. These findings demonstrate network and behavioral signatures of postpartum behavioral deficits and neurosteroid-based treatment response.

## Materials and Methods

### Animals

Adult (>P60) female Wild Type (WT) and *Gabrd*^+/−^ mice were bred and housed at Tufts University School of Medicine’s Division of Laboratory Animal Medicine facility under a 14/10 light schedule (lights on at 7:00 h) with *ad libitum* access to food and water. Littermates were used for all behavioral experiments. Females were bred with a male and the presence of a vaginal plug was used to time pregnancy (Positive plug = day 0). At Gestational Day 14 (GD14), mice were individually housed and maintained on either standard chow or SGE-516 chow (450mg/kg) which continued throughout the experiment. All procedures were approved by the Tufts University Institutional Animal Care and Use Committee and adhered to the ethical guidelines presented in the National Institutes of Health Guide for the Care and Use of Laboratory Animals. Extensive previous work from our laboratory has demonstrated behavioral deficits in *Gabrd*^−/−^ and *Gabrd*^+/−^ mice during the postpartum period [7, 16].

### Behavioral Assessments

All behavioral tests occurred between 09:00 and 16:00 h, following at least 1 h of habituation in the behavioral testing room.

#### Tail suspension

The tail suspension test was conducted as previously described [10, 16, 17]. Mice were suspended by their tail on a bar 36cm above a platform for 6 minutes. Trials were video recorded and automatically scored using an unbiased in-house Mobility App (https://github.com/researchgrant/mobility-mapper.git). Limb movement (nose, paws, and tail) were calculated from positional data automatically detected using DeepLabCut as previously described [17].

#### Maternal approach

The maternal approach test was performed as previously described [10, 16, 17]. The dams were introduced into their home cage at the opposite corner from their litter after a brief separation (10 min). The latency to approach the pups and the total time spent in contact with pups were measured as indices of maternal behavior. Only >5 s interactions were included in the time spent with the pups. Burrowing beneath or trampling the litter was not included in the measure of pup contact time. Pup calls were recorded during the 10 min period that the dam was removed from the home cage for later playback.

### LFP Recording and Analysis

Local field potential (LFP) recordings were performed in WT C57BL/6J and *Gabrd*^±/−^ mice, using in house modified prefabricated headmounts (Pinnacle 8201) as previously described [10]. The headmounts were mounted to the skull using stainless steel screws as ground, reference, and frontal cortex EEG (AP +0.75 mm, ML ± 0.3 mm, DV −2.1 mm) electrodes. One depth electrode (PFA □coated stainless □ steel wire, A□ M systems) was placed into the BLA (AP −1.35 mm, ML ± 3.45 mm, DV −5.15 mm) and was used to record the LFP. EEG and LFP recordings were acquired at 4 KHz, amplified x100, and band□ pass filtered (1□ 300 Hz, Chebyshev Type II filter). Spectral analysis was performed in python (https://github.com/SAKEverse/sake) using custom□ made scripts as previously described [10]. The power at each frequency band was normalized to the total power.

The power at each oscillation frequency was measured during predefined epochs immediately preceding the onset of pup calls (“pre-call”) and during pup-call presentation (“pup-call”). For each animal, oscillation power during each epoch was expressed as a ratio relative to the corresponding reference or baseline period. To determine whether theta-band measures could distinguish treatment groups, a binary ridge-regularized logistic regression classifier was constructed using pre-call and pup-call ratios as predictor variables and treatment condition (standard chow or SAGE) as the binary outcome. Model performance was evaluated using leave-one-animal-out cross-validation. In each fold, one animal was withheld, the model was fit using data from the remaining 15 animals, and the fitted model was used to generate a predicted probability and treatment score for the withheld animal. This procedure was repeated until every animal had served once as the held-out observation. Thus, all reported predictions were generated out of fold and were not obtained from models trained on the corresponding animal.

Cross-validated discrimination was evaluated using the receiver operating characteristic curve and its area under the curve (AUC). Classification accuracy, sensitivity, and specificity were calculated from the pooled out-of-fold predictions using a probability threshold of 0.5. For visualization, the out-of-fold treatment score for each animal was expressed as the log odds of assignment to the SAGE-treated group. Positive values therefore indicate evidence favoring SAGE treatment, whereas negative values favor standard chow treatment. Out-of-fold scores were compared between treatment groups using a two-sided between-group test. Feature contributions were summarized using the standardized ridge-regression coefficients from the final model. Coefficients were normalized to the largest absolute coefficient, such that the most influential feature had a relative value of 1.00.

### Principal Component Analysis and Multivariate Statistical Analysis

Analyses were performed using all available behavioral and electrophysiological measures. Variables included maternal approach behavior (latency to approach and total time spent with pups), stress-induced helplessness behavior (tail suspension test latency to immobility and total time immobile), and local field potential (LFP) power across oscillation frequencies. Prior to analysis, variables were centered and scaled to unit variance (z-score normalization) to account for differences in measurement units and variance.

Principal component analysis (PCA) was performed using the correlation matrix of the standardized variables. Principal components were computed by eigenvalue decomposition of the covariance matrix of the standardized data, and the proportion of variance explained by each component was calculated. Sample scores were projected onto the first two principal components (PC1 and PC2) for visualization. Ninety-five percent confidence ellipses were calculated for each treatment group from the multivariate covariance of the PCA scores.

To determine which variables contributed most strongly to separation along each principal component, PCA loading vectors were examined. Variable contributions were calculated as the squared loading of each variable normalized by the total squared loading for each principal component. To determine whether experimental groups differed in multivariate phenotype, permutational multivariate analysis of variance (PERMANOVA) was performed on Euclidean distances calculated from the standardized dataset using 9,999 permutations. Treatment effects were evaluated within *Gabrd*^*+/-*^ mice by comparing standard chow- and SAGE-treated animals. The proportion of multivariate variance explained by each factor (R^2^) and pseudo-F statistics were calculated.

To verify that significant PERMANOVA results reflected differences in multivariate centroids rather than unequal within-group variability, homogeneity of multivariate dispersion was assessed using a permutation test of distances to group centroids (PERMDISP). To identify the variables most strongly associated with treatment effects, standardized effect sizes (Cohen’s *d*) were calculated for each behavioral and electrophysiological measure. Individual variables were compared between groups using two-sided independent samples *t*-tests. To account for multiple comparisons across variables, *P*-values were adjusted using the Benjamini-Hochberg false discovery rate (FDR) procedure.

### Linear discriminant analysis

Linear discriminant analysis (LDA) was used to determine whether coordinated variation in local field potential (LFP) and behavioral measures distinguished between experimental groups. The analysis included LFP band-power measures obtained during the pre- and pup-call exposure epochs, theta-band power ratios calculated for the pre-call and pup-call periods, maternal-behavior measures, and immobility in the tail-suspension test (TST).

The LDA model was fit using the experimental group as the classification variable. Individual subjects were projected onto the first two linear discriminant axes (LD1 and LD2). The percentage of discriminant variance explained by each axis was calculated from its corresponding eigenvalue. The contribution of each variable to group separation was quantified using canonical structure coefficients, defined as the correlation between each standardized variable and the discriminant scores. Overall variable importance was calculated from the structure coefficients across the retained discriminant axes, weighted by the proportion of discriminant variance explained by each axis. Uncertainty in variable importance was estimated using 1,000 stratified bootstrap resamples that preserved the original group sizes; error bars represent bootstrap 95% confidence intervals. The signed structure coefficients for LD1 and LD2 were also examined to determine the direction of each variable’s association with the discriminant axes.

To compare contributions across measurement modalities, weighted variable-importance values were grouped into four prespecified domains: maternal behavior, TST behavior, theta ratios, and LFP band power. Values were summed within each domain and normalized to the total importance across all included variables, yielding the relative percentage contribution of each data domain to experimental-group discrimination.

### Motion Sequencing Analysis (MoSeq)

Behavioral data acquisition was performed as previously described [18]. In brief, mice were placed in an open arena (17 inch diameter and 16 inch walls and recorded for 10 mins at baseline followed by 10 mins of recording during playback of pup calls. Each mouse was tracked in 3D using a Kinect 2 (Microsoft) infra-red depth camera suspended over the arena providing a top-down view. Depth frames were retrieved at 30 frames per sec.

The data preprocessing, extraction and modeling pipeline was performed as previously described [18]. In brief, software was used to track the mouse in the enclosure and extract its position, orientation, height, size, and shape from the depth data. The 3D image of the mouse was extracted from that data using the previously described pipeline [18]. This data was used to generate the sequence of all syllables that a mouse moves through in the recording period.

To maximize power across a large set of syllable comparisons, significant changes in raw syllable usage between WT and *Gabrd*^+/−^ groups were determined using animal-level permutation testing (10,000 iterations) with Benjamini–Hochberg FDR correction (adjusted p < 0.05). Syllables significantly changed in the *Gabrd*^+/−^ mutants were then individually compared across WT, *Gabrd*^*+/-*^, and SAGE-treated animals using ANOVA with Tukey’s test for multiple comparisons. Following statistical analysis of raw syllable usage, recovery index was calculated by normalizing treated animal usage to values between the mean *Gabrd*^*+/-*^ usage (recovery index = 0) and the mean SAGE-treated usage (recovery index = 1) using the formula:

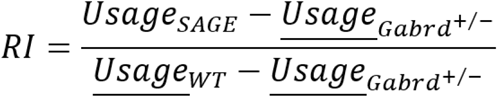

## Results

### Integrated behavioral and electrophysiological analyses identify a neural signature associated with postpartum behavioral deficits and SGE-516 treatment

To determine whether behavioral and electrophysiological measures collectively distinguished between wild type, *Gabrd*^*+/-*^, or SGE-516-treated *Gabrd*^*+/-*^ dams, we performed linear discriminant analysis (LDA) using z-scored measures of maternal approach behavior, tail suspension behavior, and local field potential (LFP) activity. Integration of these variables resulted in robust separation of wild type, *Gabrd*^*+/-*^, or SGE-516-treated *Gabrd*^*+/-*^ dams within the discriminant space (Fig. 1A). The first two discriminant axes accounted for all of the explained variance (LD1 = 65.6%; LD2 = 34.4%), indicating that a low-dimensional combination of behavioral and electrophysiological features effectively captured treatment-dependent differences.

**Figure 1.**
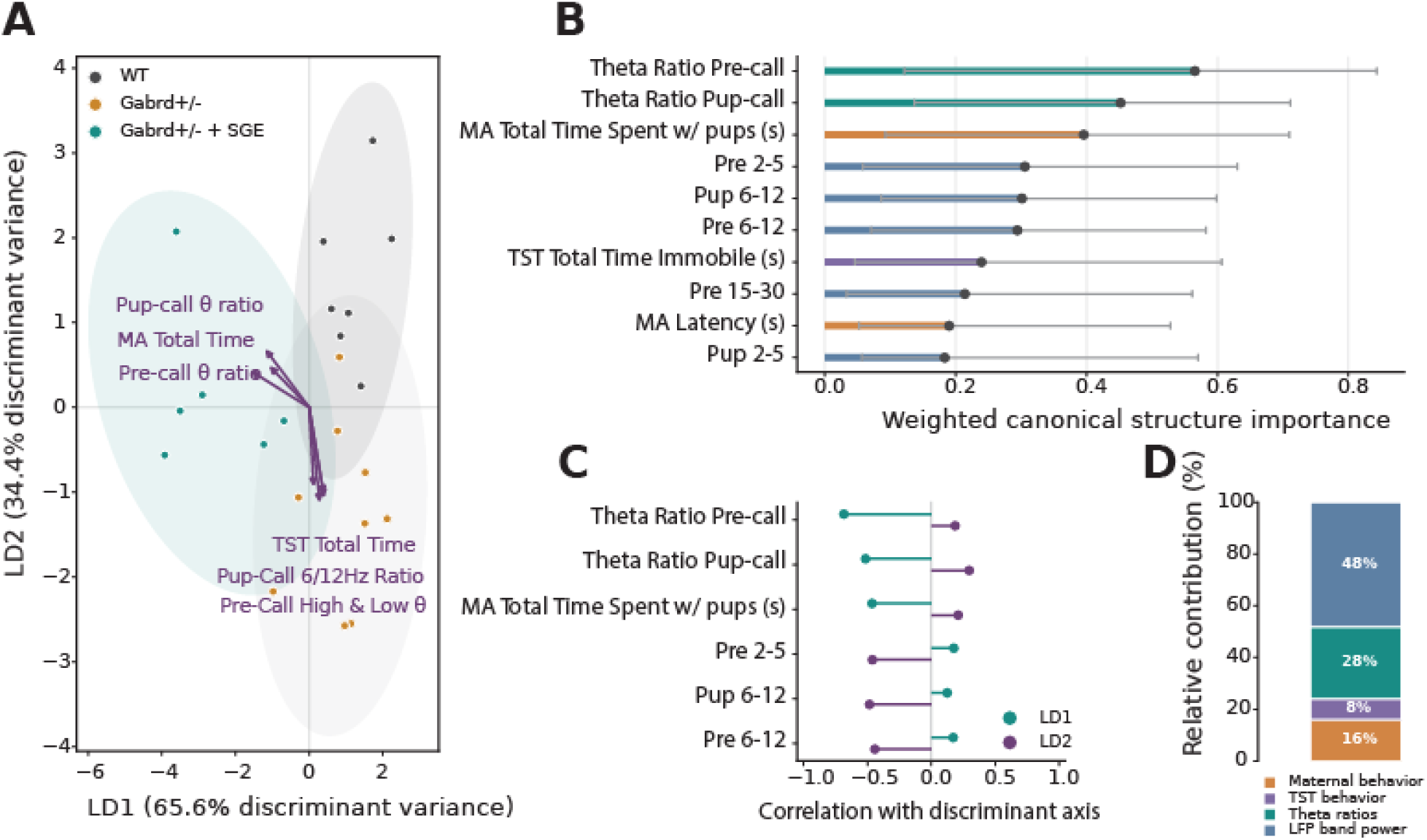
Multivariate integration of electrophysiological and behavioral measures distinguishes between genotypes and treatment. **(A)** Linear discriminant analysis (LDA) of integrated LFP and behavior variables demonstrates separation of wild type (black), *Gabrd*^*+/-*^ dams (orange), and SAGE-treated *Gabrd*^*+/-*^ dams (teal) along the first two discriminant axes (LD1 and LD2), which explain 65.6% and 34.4% of the discriminant variance, respectively. Each point represents an individual animal, and shaded ellipses indicate the 95% confidence region for each experimental group. Purple vectors represent the direction and relative contribution of individual variables to the discriminant space, with longer vectors indicating stronger contributions. **(B)** Weighted canonical structure importance for variables contributing to group discrimination. Black circles indicate the mean weighted importance of each variable, and horizontal lines represent stratified bootstrap 95% confidence intervals (1,000 resamples). Variables are ranked according to their relative contribution. Theta ratio measures before and during pup retrieval, maternal approach behavior (total time spent with pups and latency approach pups), pre-call and pup-call LFP power, and immobility in the tail suspension test (TST) were among the strongest contributors to group separation. **(C)** Canonical loadings showing the direction and magnitude of correlations between individual variables and the first (LD1, teal) and second (LD2, purple) discriminant axes. Positive and negative loadings indicate the direction of each variable’s association with the corresponding discriminant axis. **(D)** Relative contribution of each data domain to overall group discrimination. LFP band power accounted for the largest proportion of discriminative information (48%), followed by theta ratio (28%), maternal behavior (16%), and TST behavior (8%), highlighting the predominant influence of electrophysiological measures in distinguishing experimental groups. n = 8 WT, 9 *Gabrd*^*+/-*^, 7 *Gabrd*^*+/-*^ + SGE-516.

Examination of variable importance revealed that electrophysiological measures contributed most strongly to group discrimination (Fig. 1B). Theta ratio during the pre-call and pup-call periods were the highest-ranked variables, followed by maternal behavior measures, including total time spent with pups and latency to approach their pups. Canonical loading analysis demonstrated that theta-related measures and maternal behavior primarily loaded along the first discriminant axis, whereas behavioral measures associated with stress-related immobility contributed more strongly to the second axis (Fig. 1C). Consistent with these observations, partitioning variable contributions by data modality showed that LFP band power accounted for nearly half of the discriminative information (48%), followed by theta ratio (28%), maternal behavior (16%), and tail suspension behavior (8%) (Fig. 1D), indicating that electrophysiological measures were the dominant determinants of treatment separation. Wild type and *Gabrd*^*+/-*^ dams clearly separate out based on LD2, and SAGE-treated mice are pushed toward the wild type mice on that axis. LD1 reveals further loadings that could contribute to distinct physiological-behavioral state of resiliency in SAGE-treated mice.

### Theta oscillations are identified as a treatment-specific neural signature associated with SGE-516 treatment and postpartum behavioral improvement

Because theta activity emerged as the strongest contributor to group discrimination, we next examined whether theta dynamics alone could identify SGE-516-treated *Gabrd*^*+/-*^ dams. During pup-call presentation, SGE-treated *Gabrd*^*+/-*^ dams exhibited significantly greater theta ratio than standard chow-treated animals (Fig. 2A; *P* = 0.018), suggesting enhanced recruitment of theta oscillations during processing of pup vocalizations following treatment.

**Figure 2.**
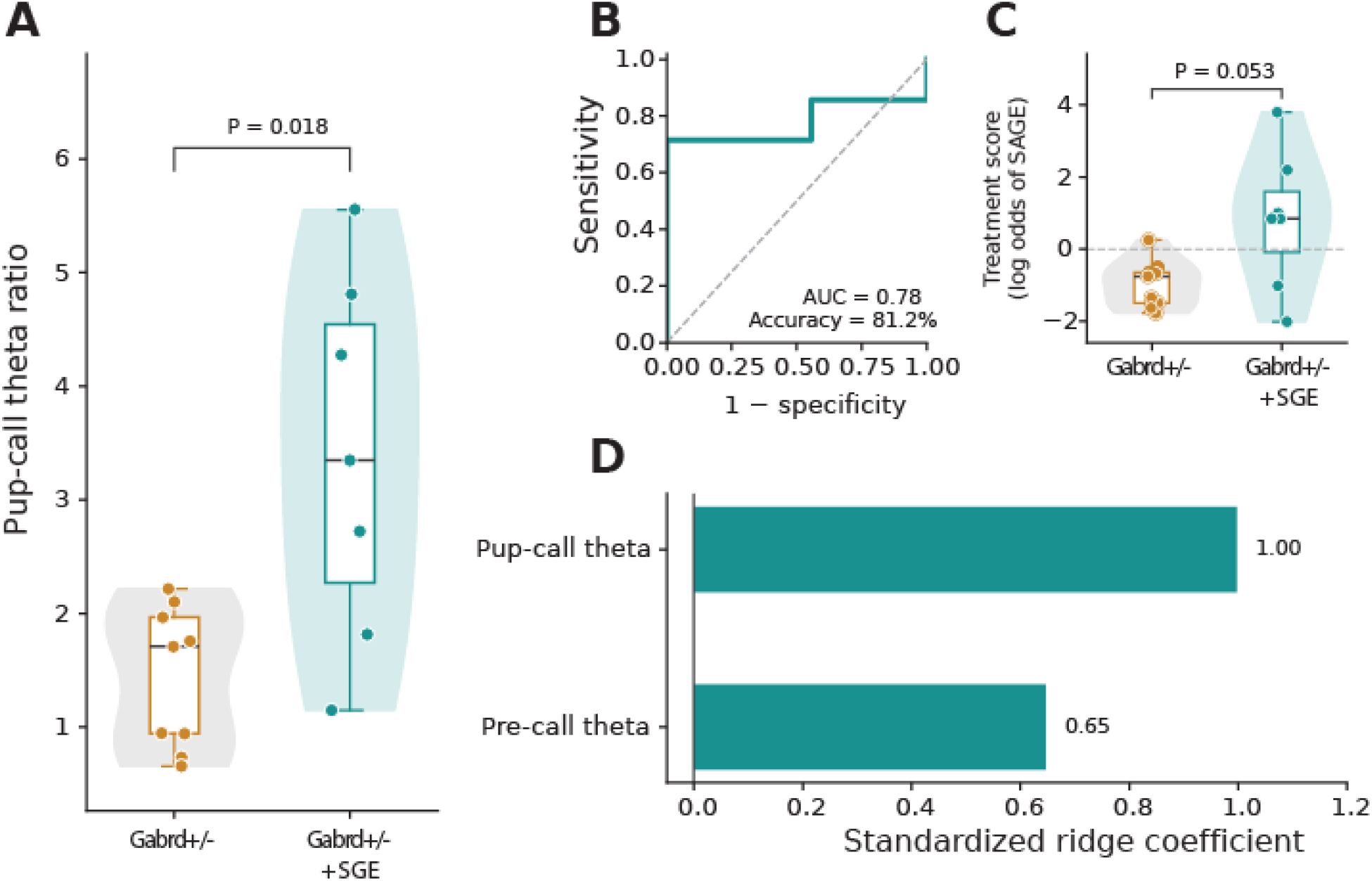
Theta activity in response to maternal cues defines an electrophysiological signature of SGE-516 treatment response. **(A)** Comparison of pup-call theta ratio between *Gabrd*^*+/-*^ dams (orange) and SGE-516-treated (teal) *Gabrd*^*+/-*^ dams. Individual animals are shown as points overlaid on violin plots with box-and-whisker summaries indicating the median and interquartile range. SGE-516 treatment significantly increased pup-call theta activity compared with standard chow-treated *Gabrd*^*+/-*^ dams (*P* = 0.018). **(B)** Performance of a binary ridge logistic regression classifier trained using pre-call and pup-call theta ratios to distinguish SGE-516 from standard chow treatment. Classifier performance was evaluated using leave-one-animal-out cross-validation (LOOCV). The receiver operating characteristic (ROC) curve yielded an area under the curve (AUC) of 0.78 and an overall classification accuracy of 81.2%, demonstrating good discrimination between treatment groups. **(C)** Out-of-fold treatment scores generated during cross-validation for individual animals, expressed as the log odds of SGE-516 treatment classification. Positive values indicate a greater likelihood of classification as SGE-516-treated, whereas negative values indicate classification as *Gabrd*^*+/-*^ dams. SGE-516-treated *Gabrd*^*+/-*^ dams exhibited higher treatment scores than standard chow-treated animals, with a trend toward significance (*P* = 0.053). **(D)** Standardized ridge regression coefficients identifying the electrophysiological features contributing to treatment classification. Pup-call theta ratio was the strongest predictor of SGE-516 treatment (standardized coefficient = 1.00), followed by pre-call theta ratio (0.65), indicating that enhanced theta activity during pup vocalizations is the dominant electrophysiological feature associated with SGE-516 treatment. n = 9 *Gabrd*^*+/-*^, 7 *Gabrd*^*+/-*^ + SGE-516.

To determine whether these electrophysiological features were sufficient to predict treatment assignment, we trained a ridge logistic regression classifier using only pre-call and pup-call theta ratios. Using leave-one-animal-out cross-validation, the classifier correctly classified 13 of 16 animals (81.2% accuracy), achieving an area under the receiver operating characteristic curve (AUC) of 0.78, with a sensitivity of 71.4% and specificity of 88.9% (Fig. 2B). The classifier produced higher out-of-fold treatment scores for SGE-516-treated *Gabrd*^*+/-*^dams than standard chow-treated animals, with a strong trend toward significance (*P* = 0.053; Fig. 2C). Examination of standardized model coefficients revealed that pup-call theta ratio was the strongest predictor of treatment status, contributing approximately 1.5-fold more than pre-call theta ratio (Fig. 2D). Together, these findings identify enhanced theta activity during pup vocalizations as the principal electrophysiological feature associated with SGE-516 treatment and demonstrate that a simple theta-based model accurately distinguishes treated animals from controls.

Given that theta oscillations and maternal care were identified as drivers of group differences between wild type, *Gabrd*^*+/-*^, or SGE-516-treated *Gabrd*^*+/-*^ dams, we analyzed the relationship between these two variables. While there is no significant differences in the theta ratio (2-5/6-12Hz) between wild type, *Gabrd*^*+/-*^, or SGE-516-treated *Gabrd*^*+/-*^ dams pre-call, there is a trend towards a reduction in *Gabrd*^*+/-*^ dams which is restored with SGE-516 treatment (Figure 3A). Further, there is a positive correlation between the ratio of the power of 2-5Hz to 6-12Hz oscillations in the BLA at baseline (pre-call) where a higher low:high theta ratio is associated with an increased amount of time spent with the pups during the maternal approach test (Figure 3B). These findings point to an interaction between BLA network states and postpartum behaviors.

**Figure 3.**
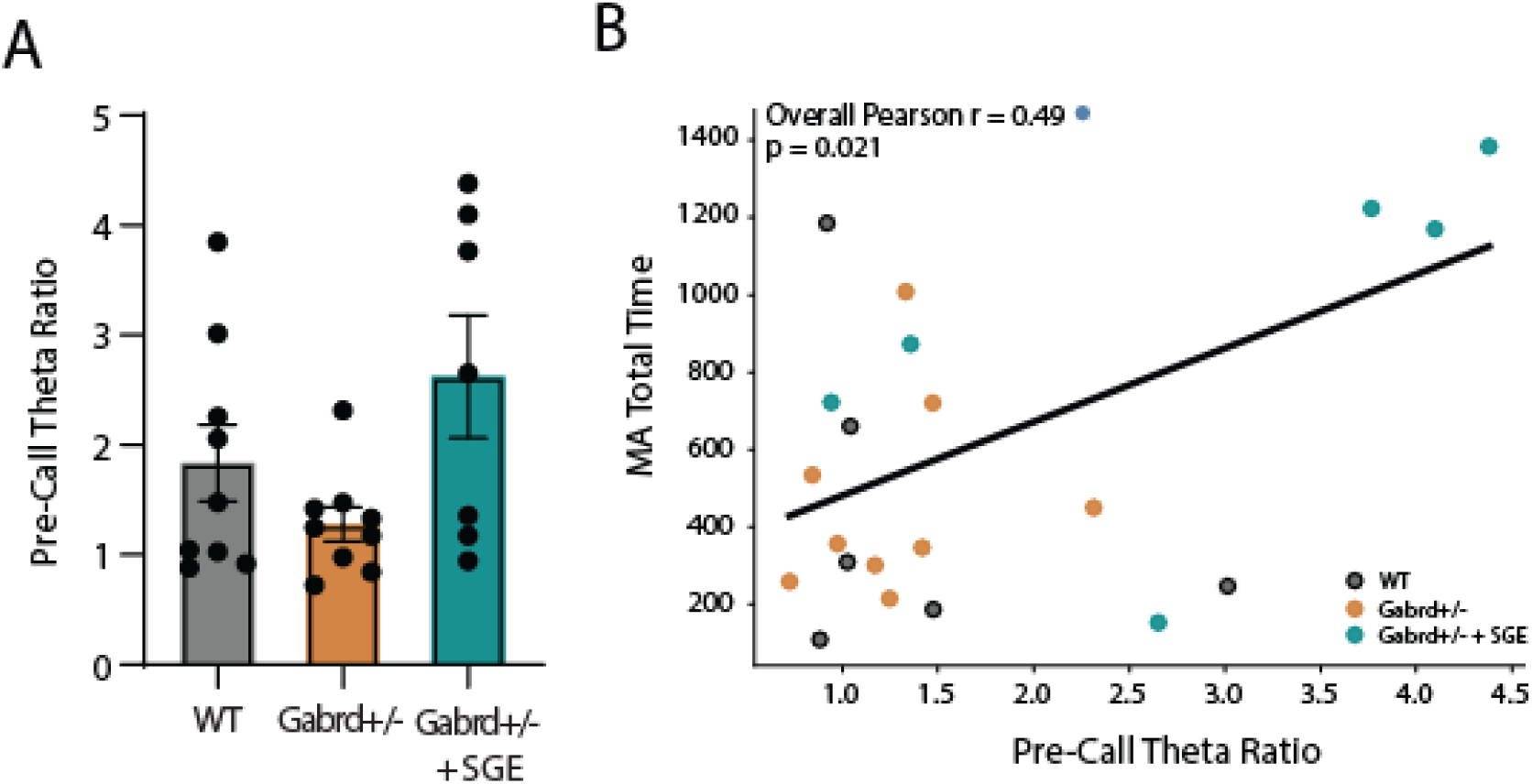
Theta activity correlates with maternal behavior. **(A)** The average ratio of the power of 2-5/6-12 Hz oscillations in the BLA at baseline (pre-call) in wild type, *Gabrd*^*+/-*^ dams, and SGE-516-treated *Gabrd*^*+/-*^ dams. **(B)** A linear regression analysis of the pre-call ratio of the power of 2-5/6-12 Hz oscillations relative to total time spent with their pups during the maternal approach test demonstrates a positive correlation between pre-call theta ratio and amount of time spent with their pups. n = 7-9 dams per group. n = 8 WT, 9 *Gabrd*^*+/-*^, 7 *Gabrd*^*+/-*^ + SGE-516.

### Unique behavioral syllables identified in Gabrd^+/-^ dams are restored with SGE-516 treatment

To expand the behavior analysis beyond traditional human-defined end-points, we performed a non-biased analysis of sub-second naturalistic movements (behavioral syllables) using Motion Sequencing analysis (MoSeq) to reveal the full structure of behavior during pre-call and pup-call playback (Figure 4A). Initial characterization of the *Gabrd*^*+/-*^ phenotype revealed 12 syllables (Figure 4B) significantly changed from wild type during pup-call playback (FDR adjusted p < 0.05, see supplementary Figure 4A for all 70 syllables analyzed). These 12 syllables were used to construct a disrupted postpartum behavioral phenotype to assess the degree of recovery toward the wild type phenotype in SAGE treated animals. Of the 12 syllables, 3 syllables showed a significant change from *Gabrd*^*+/-*^ and no significant difference from wild type (Figure 4C), indicating full recovery. 6 syllables showed a trend toward recovery, but no significant change from *Gabrd*^*+/-*^ (Figure 4D). While 3 syllables showed no improvement from *Gabrd*^*+/-*^ (Figure 4E), it should also be noted that none of syllables in the *Gabrd*^*+/-*^ phenotype exhibited a significantly worse recovery index due to SAGE-treatment (Figure 4F). These findings clearly corroborate previous work showing that SGE-516 treatment has a restorative behavioral effect in preclinical models exhibiting postpartum behavioral deficits [16], but they also reveal an incomplete restoration of the integrated behavior structure of the wild type phenotype.

**Figure 4.**
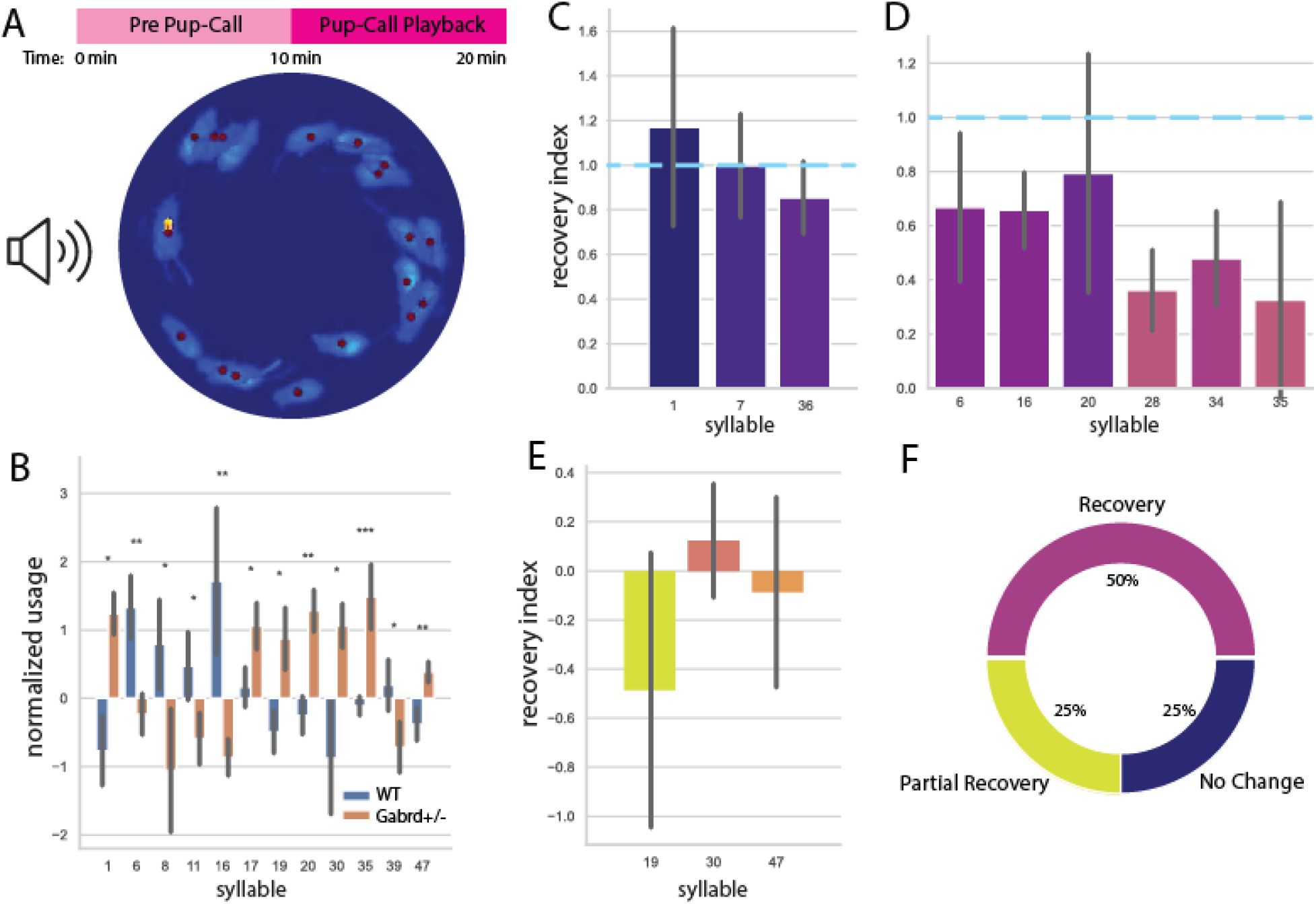
SGE-516 treatment improves postpartum behavioral deficits. **(A)** MoSeq recording timeline and representative overlay of several examples of a single behavioral syllable. **(B)** Analysis of syllable usage shows 12 syllables exhibiting a significant change between wild type (WT) and *Gabrd*^*+/-*^dams during pup-call playback, normalized by animal to pre-call usage. **(C-E)**. Recovery index calculated by normalizing SGE-516-treated animals syllable usage values between WT (RI=1) and *Gabrd*^*+/-*^ (RI=0), with panels grouped by a significant recovery (C), partial recovery (D) and no recovery from *Gabrd*^*+/-*^ phenotype. **(F)** Summary of proportions of syllables affected SGE-516-treated animals. n = 12 WT, 11 *Gabrd*^*+/-*^, 11 *Gabrd*^*+/-*^ + SGE-516.

## Discussion

Behavioral deficits restricted to the postpartum period have been previously reported in *Gabrd*^*-/-*^ and *Gabrd*^*+/-*^ dams [7, 16], with reported deficits in maternal care and increased pup mortality in both *Gabrd*^*-/-*^ and *Gabrd*^*+/-*^ dams [7] which were improved with GABA PAM treatments [7, 16], including SGE-516 [16]. However, these approaches largely relied on a standard behavioral battery with controverted translational relevance. Here we expanded the interrogation of the postpartum phenotype in *Gabrd*^*+/-*^ dams focusing on comprehensive and unbiased behavioral analyses. Here we demonstrate genotype differences in the behavioral profiles of *Gabrd*^*+/-*^ dams using unsupervised analytical approaches. PCA and LDA analyses identified a neural signature associated with postpartum behavioral deficits and SGE-516 treatment response. Further, the unbiased MoSeq behavioral screening identified genotype differences in behavioral responses to pup calls and a recovery of behavior with SGE-516 treatment. These multivariable and unbiased behavioral analyses clearly demonstrate behavioral abnormalities in *Gabrd*^*+/-*^ dams compared to wild type and the ability of SGE-516 to improve behavioral outcomes. These approaches may be useful for screening antidepressant compounds for effectiveness without the caveat of examining subjective behavioral outcomes.

Previous work from our laboratory also demonstrated a role for theta oscillations in the BLA in emotional processing, including the behavioral expression of fear and stress-induced helplessness [10, 15]. We demonstrated that chronic stress exposure alters low:high theta oscillations in the BLA which correlates with the behavioral expression of fear [15]and stress-induced helplessness [10]. Here we examined whether oscillations in the BLA may also relate to postpartum behaviors. We demonstrate that the separation between wild type and *Gabrd*^*+/-*^ dams is primarily driven by oscillations in the BLA, particularly oscillations in the theta range. Consistent with our previous work demonstrating that allopregnanolone treatment was capable of modulating oscillations in the BLA and restoring the healthy network state following chronic stress exposure [10], here we demonstrate that the effects of SGE-516 treatment are largely driven by the ability to alter theta oscillations in the BLA. Together, these findings suggest a neural signature associated with behavioral deficits in postpartum *Gabrd*^*+/-*^ dams is restored with SGE-516 treatment, suggesting a neural signature of SGE-516 treatment response.

The fact that there is a relationship between 2-5/6-12Hz and maternal care suggests a relationship between the BLA network states and postpartum behaviors. The 2-5/6-12Hz ratio has previously been associated with stress and threat responding [10, 14, 15]. Interestingly, an increased 2-5/6-12Hz ratio is associated with better maternal care, which suggests that this may be a signature of the salience of the stimulus. We demonstrate that SGE-516 increases the 2-5/6-12Hz ratio in *Gabrd*^*+/-*^ dams, perhaps improving the neural and behavioral response to pup cues. This is unique from previous findings in male mice demonstrating that following chronic stress, the ability of SGE-516 to reduce 2-5/6-12Hz oscillations in the BLA was correlated with behavioral improvement in stress-induced helplessness tests. Thus, it appears that these neural signatures are context dependent and may involve salient cues that require an adaptive response.

The data presented here demonstrate altered neural and behavioral responses to pup calls in *Gabrd*^*+/-*^ dams which is reminiscent of the altered responses of women with postpartum depression to hearing their infants cry [19]. Functional neuroimaging studies demonstrate that hearing an infant cry engages a distributed maternal caregiving network, with the amygdala serving as an important component linking the detection of emotionally salient infant cues with motivational and behavioral responses. In postpartum women, an infant’s cry, in particular the cry of one’s own infant, recruits the amygdala together with the insula, anterior cingulate and prefrontal cortices, striatum/nucleus accumbens, hypothalamus, thalamus, and sensorimotor regions [19, 20]. Importantly, the maternal response to infant distress appears to involve not simply changes in amygdala activity but alterations in the functional integration of the amygdala with broader cortical and subcortical networks. For example, amygdala connectivity with regions involved in motor preparation, prefrontal regulation, and motivational processing is modulated during exposure to infant cry, consistent with a mechanism through which detection of infant distress is translated into an organized caregiving response [20, 21]. Depressive symptoms are associated with altered extended-amygdala connectivity with the nucleus accumbens and dorsomedial prefrontal cortex during processing of infant distress, while other studies demonstrate reduced recruitment of prefrontal, cingulate, striatal, and thalamic regions in women with postpartum depression responding to their own infant’s cry [19, 20]. Moreover, greater amygdala responses to an infant’s cry have been associated with greater subsequent maternal sensitivity, suggesting that adaptive maternal responding depends on coordinated activity across salience, emotion-regulation, motivational, and action-preparation networks rather than amygdala activation in isolation [20, 22]. These findings are consistent with the altered oscillations in the BLA in *Gabrd*^*+/-*^ dams in response to hearing playback of their pup calls.

The PPD-like integrated behavior phenotype identified by MoSeq analysis shows a more complete picture of the degree of recovery due to SGE-516 treatment. While most syllables showed either robust or moderate recovery, a few remained unchanged. It is unclear if the unaffected syllables are translationally relevant to PPD. Further optimization of recovery of the PPD-like syllables could be a valuable tool in screening new treatment schedules or drug analogues. Future treatment development would also benefit from mitigating the drug-induced effects on syllables outside of the PPD-like set, in order to reduce unwanted side effects.

These findings identify a neural signature of treatment response which may serve as a useful biomarker in clinical populations. EEG represents a promising, scalable approach for identifying neurophysiological biomarkers of depression and predicting treatment response. In major depressive disorder (MDD), abnormalities have been reported across multiple EEG domains, including frontal alpha asymmetry, altered alpha and theta power, disrupted functional connectivity and coherence, sleep-related measures, and event-related potentials, although the reproducibility of individual diagnostic markers remains variable [13]. Among these measures, frontocentral theta activity and particularly theta activity localized to the rostral anterior cingulate cortex (rACC) have emerged as some of the more consistently replicated predictors of antidepressant response, with greater pretreatment rACC theta current density being associated with subsequent response to monoaminergic antidepressants [23, 24]. Machine-learning approaches integrating spectral power, asymmetry, connectivity, and clinical features further support the potential of multidimensional EEG signatures for predicting antidepressant response, although methodological heterogeneity and limited external validation currently restrict clinical implementation. In contrast to MDD, PPD EEG biomarkers have not been extensively explored. This represents an important opportunity because PPD is associated with altered activity and connectivity within prefrontal, limbic, salience, and reward networks, including disrupted medial prefrontal–ventral striatal connectivity, suggesting that EEG measures of oscillatory activity and functional network organization could provide accessible physiological correlates of these circuit abnormalities. This hypothesis is supported by findings in the current manuscript. Collectively, these findings support the development of neural signatures to identify biologically distinct subgroups of women with PPD and, importantly, to predict treatment response.

## Data Availability

Data will be made available upon request.

## Acknowledgments

The authors would like to thank the Datta lab for the development of the MoSeq approach and for facilitating the dissemination and adoption of this approach. The authors would also like to thank Sage Therapeutics for supplying the SGE-516 chow.

## Author Contributions

L.H. performed all of the behavioral assessments, performed analysis and interpretation of the data, and contributed to writing the manuscript. G.W. established the MoSeq behavioral approach, performed all the MoSeq analysis and interpretation of the data, and contributed to writing the manuscript. K.D. performed all the LFP acquisition and contributed to the analysis and interpretation of the data, and contributed to writing the manuscript. J.M. performed the PCA and LDA analysis, contributed to the interpretation of the data, and contributed to writing the manuscript.

